# Electrophysiological recording of human neuronal networks during suborbital spaceflight

**DOI:** 10.1101/2022.10.25.512608

**Authors:** Andie E. Padilla, Candice Hovell, Jeremy Mares, Veerle Reumers, Binata Joddar

## Abstract

The use of microfluidic tissue-on-a-chip devices in conjunction with electrophysiology (EPHYS) techniques has become prominent in recent years to study cell-cell interactions critical to the understanding of cellular function in extreme environments, including spaceflight and microgravity. Current techniques are confined to invasive whole-cell recording at intermittent time points during spaceflight, limiting data acquisition and overall reduced insight on cell behaviour. Currently, there exists no validated technology that offers continuous EPHYS recording and monitoring in physiological systems exposed to microgravity. In collaboration with imec and SpaceTango, we have developed an enclosed, automated research platform that enables continuous monitoring of electrically active human cell cultures during spaceflight. The Neuropixels probe system (imec) will be integrated for the first time within an engineered in-vitro neuronal tissue-on-a-chip model that facilitates the EPHYS recording of cells in response to extracellular electrical activity in the assembled neuronal tissue platform. Our goal is to study the EPHYS recordings and understand how exposure to microgravity affects cellular interaction within human tissue-on-a-chip systems in comparison to systems maintained under Earth’s gravity. Results may be useful for dissecting the complexity of signals obtained from other tissue systems, such as cardiac or gastrointestinal, when exposed to microgravity. This study will yield valuable knowledge regarding physiological changes in human tissue-on-a-chip models due to spaceflight, as well as validate the use of this type of platform for more advanced research critical in potential human endeavours to space.

## 1. Introduction

Understanding the EPHYS characteristics of neuronal cellular networks is essential to comprehending the mechanisms by which they function. Once limited to whole cell recording in 2D arrays, advancements in tissue engineering and MEA technologies have made possible the extracellular characterization of networks as a whole in 3D. From this knowledge, it may be possible to pinpoint critical events which shed light on the behaviour of cultures in an endless array of models from disease pathogenesis and drug toxicity screening to extreme environmental studies. This study aims to utilize these state-of-the-art tools to identify the effects of microgravity on human iPSC derived glutamatergic neurons in efforts to translate this to current knowledge regarding potential human endeavours to space.

### 1.1 Neuronal Electrophysiology and Micro-electrode arrays

EPHYS techniques are well understood as a validation method for in vitro models. They operate by recording action potentials induced by the inward and outward flux of Na^+^, Ca^+^, K^+^, and Cl^-^ ions through the lipid membrane. Early work relied on single or whole-cell recording via a method known as patch-clamp. This technique involves isolating mechanosensitive ion channels on the cell membrane by using the suction from a micropipette to create a seal along a small portion of the cell membrane [1]. Ion response to stimulation via drug delivery or induced current is then measured in terms of voltage output via an electrode in the micropipette. Though this technique provides a well-established and reliable method for obtaining EPHYS data, it is bounded to a single-cell’s response.

Developments in this field have progressed towards the use of micro-electrode arrays (MEAs) for EPHYS data acquisition. These devices consist of multiple electrodes which can measure action potentials from numerous cells during a single acquisition. The extracellular nature of this technique maintains the viability of cultures over long periods. This proves to be especially useful for models requiring long-term analysis [2,3]. The incorporation of 3D tissue models into MEAs has also produced a wide array of electrode designs optimized to suit the needs of specific models such as mesh, probe, and plate types. These developments allow for expanded spatial resolution recording and high-throughput data acquisition which provides insight into network function as a whole.

In this work, we validate a state-of-the-art MEA, developed by imec USA, known as the neuropixels probe for use in in vitro models. This device features 960 low impedance titanium nitride recording sites connected to 384 parallel, configurable, low noise recording channels along a 10mm shank [4]. Though its current use is largely for in vivo studies, the streamlined design and accompanying data acquisition technology prove it to be suitable for use in in vitro models.

## 2. Material and methods

### 2.1 Ostemer device fabrication

3D printed molds (Accura 5330) of the device were obtained from Protolabs (Maple Plain, MN, USA). These were used to cast silicone molds from Sylgard™ 184. From this, final devices are cast from Ostemer® 322 following the recommended protocol. After the completion of the UV curing step, the device is placed onto a microscope slide with a Neuropixels probe set in place, then oven-cured for 1 hour at 90°C.

### 2.2 Cell culture and seeding

Human glutamatergic neurons derived from iPSCs (iCell® GlutaNeurons) were purchased from Cellular Dynamics International (Madison, WI, USA). Cells were thawed and resuspended in 9 mL complete BrainPhys maintenance medium (BrainPhys Neuronal Medium, iCell DopaNeurons Supplement, iCell Nervous System Supplement, N-2 Supplement, Laminin, and Penicillin-Streptomycin). Cells were centrifuged at 400 rpm for 5 minutes and resuspended in 500µL complete maintenance medium. 150µL of this suspension was then taken and placed into a 1.5mL Eppendorf tube and centrifuged again. Cells were then resuspended in 5µL complete maintenance medium. 24µL Matrigel was added to the suspension and gently mixed by pipetting to achieve a density of 300,000 cells/24µL Matrigel. The gel was pipetted into the device gel chamber and incubated for 30 minutes at 37°C, 5% CO2. Complete maintenance medium was then added to the device and exchanged every 1-2 days.

### 2.3 Sample preparation and integration for suborbital flight

Four biological replicates were prepared as previously described in preparation for the suborbital flight, two samples would be integrated into a CubeLab developed by SpaceTango and two remained on the ground to be used as ground controls. Similarly, two devices seeded with blank Matrigel would be sent on the flight to be used as positive controls. During culture, luer connector components (McMaster Carr #51525K211) served as hydrostatic reservoirs. Upon integration into the CubeLab, these components were replaced with alternate connectors (McMaster Carr #5463K36) fitted with tubing to accommodate for pumps which will facilitate automated media exchange during the flight

### 2.4 EPHYS measurements

Electrophysiology measurements were obtained using a Neuropixels probe (imec, Leuven, Belgium) integrated into the device. Data acquisition was performed using a PXIe-1071 (National Instruments, Austin, TX, USA) fitted with a neuropixels card as recommended by imec.

### 2.5 Device compatibility testing

A live/dead assay was performed to assess cell viability and overall device biocompatibility for the Ostemer device. Non-differentiated SH-SY5Y cells were cultured for 7 days in 1.5% gelatin-alginate hydrogel seeded in the Ostemer device. Media exchange was performed every 1-2 days. Live/Dead assay was performed using a kit (Invitrogen, Gibco, CA, USA) and used according to manufacturer’s protocol. Images were processed using ImageJ to quantify percent viability of cells studied.

### 2.6 Immunocytochemistry

iCell® GlutaNeurons in Matrigel were seeded and maintained in an Ostemer device at a density of 300,000 cells/24µL. Immunocytochemistry was performed via DAPI and MAP-2 immunostaining after 21 days to assess robust network formation and growth within the device.

## 3. Theory and calculation

### 3.1 Device Design

Microfluidic devices extend the ability to model and analyse the desired culture within a single platform. For the purposes of this study, the device design was intended to accommodate for the neuropixels probe, cell culture, and culture medium.

Initial testing was conducted using an open loop model (Fig. 1A). The design was later modified and optimized (Fig. 1B) to promote media circulation and prevent the loss of media from the microfluidic circuit. The optimized design accommodates 100 µL of media, supplemented every 1-2 days, and 24 µL of hydrogel. The port size was also reduced to allow for the incorporation of a microfluidic circuit performing autonomous media exchange (Fig. 1C). Automation granted an additional 2mL of media, reserved outside of the chip, ensuring tissue survival for several days for transport to and from the launch site and in the event of a delay. The samples and circuitry were housed in a CubeLab developed by SpaceTango which also provided a closed, sterile environment, 37°C incubation, and a 5% CO2 atmosphere.

**Figure 1:**
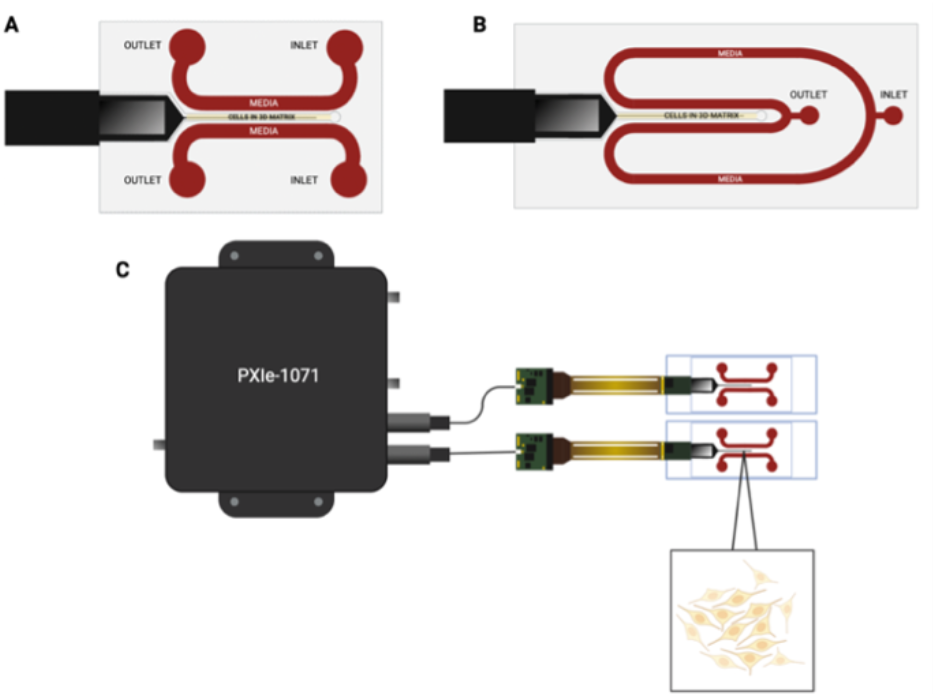
(A) and (B) show the initial and selected design models for the microfluidic device. Both feature a 24 µL hydrogel chamber and probe slot. Design B was developed and selected to accommodate for closed-loop automated media exchange during the flight. (C) Shows the in-flight data acquisition setup.

## 4. Results

Representative images collected from different areas in the device (Fig, 2A) showed that the device was biocompatible. This was confirmed via live/dead assay which show approximately 90% viability after 7 days in vitro across all regions, with the calculated viability being 92.5%, 89.5%, and 90.5% for banks 0,1, and 2 respectively (Fig. 2B).

**Figure 2:**
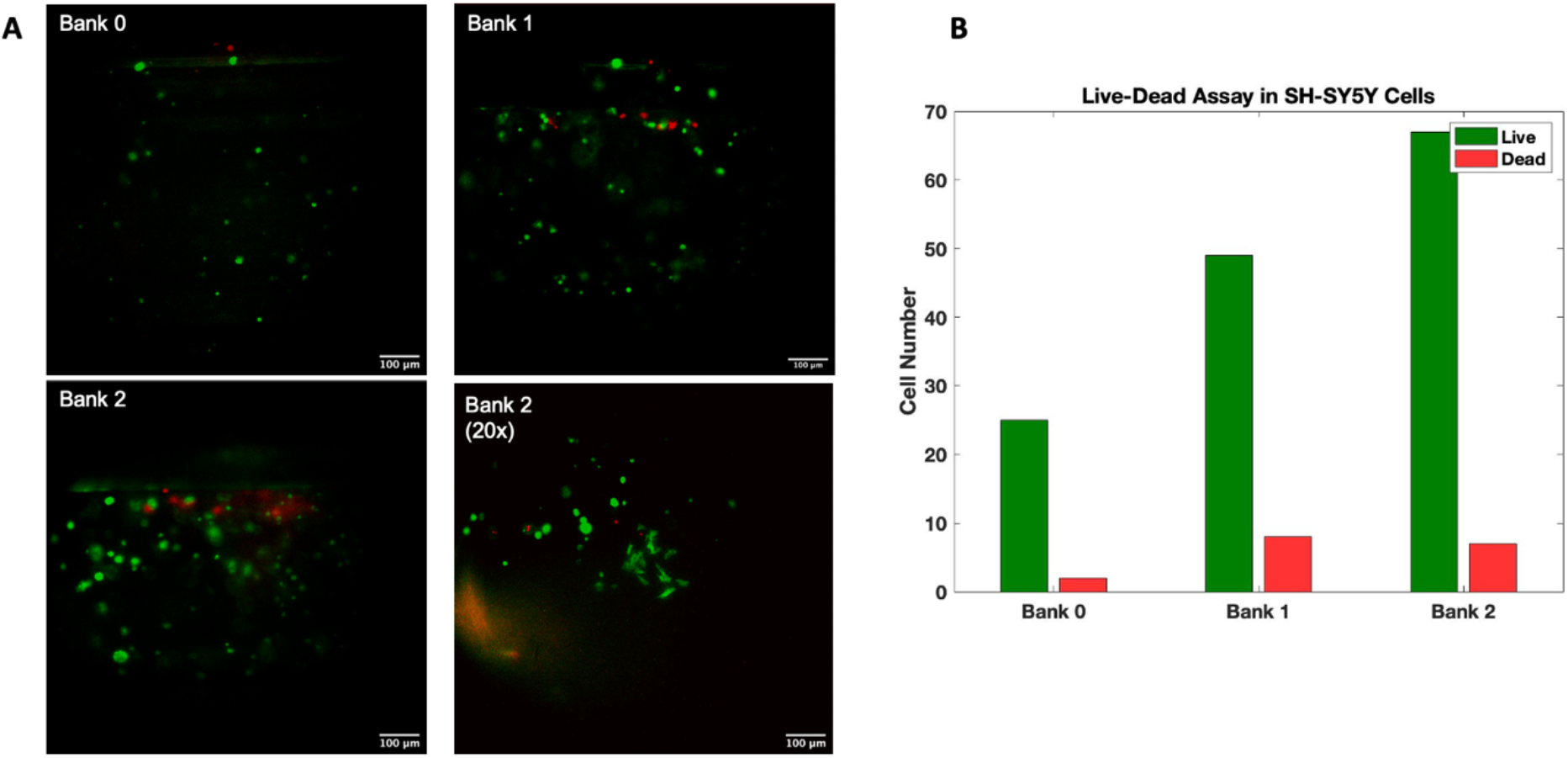
Live/Dead assay was performed on differentiated SH-SY5Y seeded in 1.5% gelatin-alginate hydrogel in ostemer device after 7 days across three regions of the device dictated by probe placement. Results show approximately cell viability ≥ 90% across all regions of the device in terms of viable (green) cells.

Immunostaining with MAP2 and Counterstaining with DAPI show the formation of robust neuronal networks in the device after a 3-week culture period in the device (Fig. 3). Figures 3A and C show the presence of cells on and around the probe. Clustering of the cells around the probe was observed suggesting preferential cell alignment in these areas in comparison to other regions of the device.

**Figure 3:**
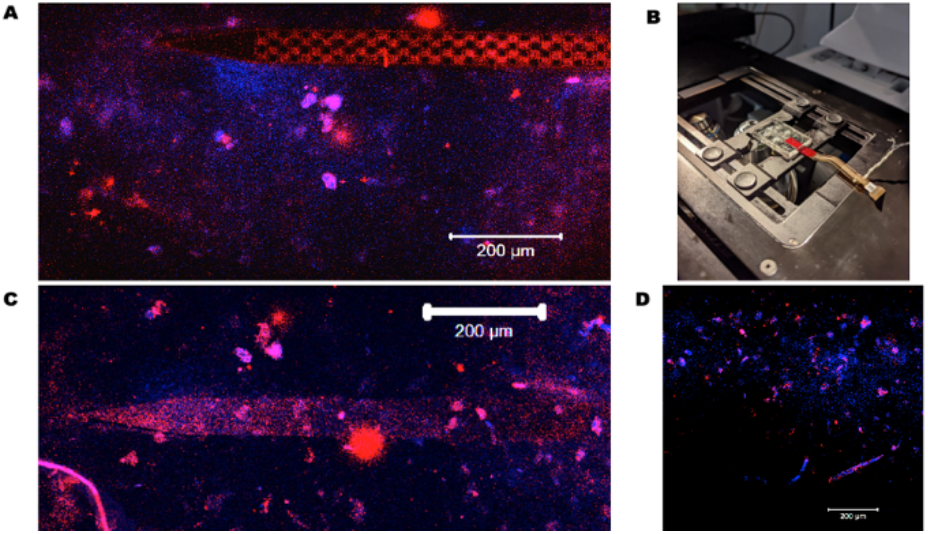
(A) and (C) show the probe with the Glutamatergic neurons networked in the Matrigel. The cells were stained using antibodies for the neuronal marker MAP-II (red) and counterstained using the nuclear stain DAPI (blue). The representative image (A) was acquired using the set-up depicted in (B) where the sample was placed (face-down) for imaging. In (C) the sample was placed face-up for imaging. Shown in (D) is a z-stacked projection image to depict the cell density and neuronal networks developed after a 3-week culture period in the device.

Biological samples were prepared and seeded 8 days in advance of flight. EPHYS signals were observed after 2 days in culture. Fig. 4 shows representative images of spontaneous neuronal firing during a 1-minute acquisition from two samples on day 4. Data from this acquisition was plotted in 0.1 second intervals to show individual peaks more clearly. Both samples exhibit strong firing activity, which suggests healthy network formation and intercellular communication.

**Figure 4:**
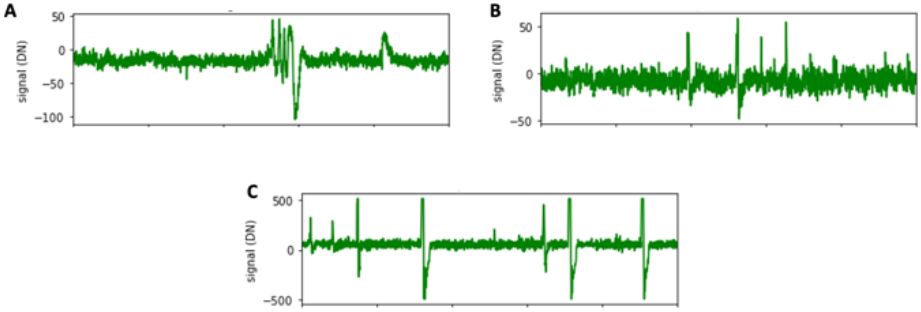
EPHYS recording of two tissue samples after 4 days in culture. (A) and (B) show acquisitions from sample 1, while (C) reflects sample 2.

## 5. Discussion and Conclusions

Live/dead assay and microscopy data indicate that cell viability is well maintained within the device. This provides confidence that EPHYS data acquired from this model is produced from healthy neuronal networks, which is critical to achieve an accurate and complete understanding of the effects of microgravity following a suborbital flight. Preliminary data acquisition from the devices shows spontaneous neuronal firing activity after 4 days in culture, validating the use of neuropixel technology in tissue engineering applications. As per our research, there exists no other validated hardware for the EPHYS recording of human neuronal networks in microgravity. Successful completion of a suborbital flight experiment with this integrated EPHYS neuronal tissue on a chip model will enable others in the field to adopt this technology for spaceflight studies.

## Acronyms/Abbreviations

(EPHYS): Electrophysiology
(iPSC): induced pluripotent stem cells
(2D): two-dimensional
(3D): three-dimensional
(MEA): microelectrode arrays

## Acknowledgements

The authors would like to acknowledge the NASA grant ‘Electrophysiology recording of neuronal networks during suborbital spaceflight’ in collaboration with imec.

